# Quantification of nuclear transport inhibition by SARS-CoV-2 ORF6 using a broadly applicable live-cell dose-response pipeline

**DOI:** 10.1101/2021.12.10.472151

**Authors:** Tae Yeon Yoo, Timothy Mitchison

## Abstract

SARS coronavirus ORF6 inhibits the classical nuclear import pathway to antagonize host antiviral responses. Several models were proposed to explain its inhibitory function, but quantitative measurement is needed for model evaluation and refinement. We report a broadly applicable live-cell method for calibrated dose-response characterization of the nuclear transport alteration by a protein of interest. Using this method, we found that SARS-CoV-2 ORF6 is ~15 times more potent than SARS-CoV-1 ORF6 in inhibiting bidirectional nuclear transport, due to differences in the NUP98-binding C-terminal region that is required for the inhibition. The N-terminal region promotes membrane binding and was required for activity, but could be replaced by constructs which forced oligomerization in solution. Based on these data, we propose that the hydrophobic N-terminal region drives oligomerization of ORF6 to multivalently cross-link the FG domains of NUP98 at the nuclear pore complex, and this multivalent binding inhibits bidirectional transport.

## Introduction

Macromolecular transport across the nuclear envelope occurs through nuclear pore complexes (NPCs) and is tightly regulated in eukaryotic systems. The transport channel of the NPC is densely filled with intrinsically disordered regions of the FG-nucleoporins (FG-NUPs) enriched in phenylalanine-glycine repeats (Denning et al., 2003; Kim et al., 2018; Lin and Hoelz, 2019). The FG-repeat regions constitute a permeability barrier that specifically interacts with soluble transport carrier proteins, including importin-beta family proteins, to selectively facilitate their diffusion through the pore along with cargoes bound to them (Feldherr and Akin, 1997; Fried and Kutay, 2003; Keminer and Peters, 1999; Ribbeck and Gorlich, 2001). The classical nuclear import pathway involves importin beta-1 (KPNB1) as a transport carrier and a class of adaptor proteins called importin alphas (KPNAs) which bind classical nuclear localization signals (NLSs) in complex with KPNB1 (Lange et al., 2007). The classical nuclear export pathway is mediated by exportin-1 (CRM1/XPO1), another transport carrier in the importin-beta family, which directly binds to leucine-rich classical nuclear export signals (Stade et al., 1997). Directionality of cargo transport is provided by compartment-specific assembly and disassembly of cargo-carrier complexes, which is mediated by the small GTPase RAN (Christie et al., 2016; Macara, 2001; Matsuura, 2016).

Nucleocytoplasmic transport is required for several innate immune anti-viral pathways; for example, the interferon (IFN) response requires nuclear import of phosphorylated STAT1 (Ivashkiv and Donlin, 2014). SARS coronaviruses, like several other viruses, were reported to inhibit the nucleocytoplasmic transport system of host cells during infection, resulting in reduced innate immune signaling (Lei et al., 2020; Miorin et al., 2020; Sajidah et al., 2021; Tessier et al., 2019; Xia et al., 2020; Yarbrough et al., 2014). A small protein that is unique to these viruses, Open Reading Frame 6 (ORF6), was shown to inhibit the KPNB1-mediated classical nuclear import pathway (Kimura et al., 2021). Several inhibition mechanisms were proposed. Early studies on SARS-CoV-1 proposed that its ORF6 tethers KPNA2 to membranes of the endoplasmic reticulum (ER) and Golgi apparatus to sequester KPNB1 (Frieman et al., 2007). Interaction of SARS-CoV-2 ORF6 with KPNA2 was also demonstrated (Miorin *et al*., 2020; Xia *et al*., 2020). Recent studies on SARS-CoV-2 ORF6 identified NUP98-RAE1 complex, an NPC component possessing a long FG-repeat domain, as a new binding partner of ORF6 and implicated this interaction in its inhibitory function (Addetia et al., 2021; Gordon et al., 2020; Miorin *et al*., 2020). Miorin et al. proposed that ORF6 binding to the NUP98-RAE1 complex inhibits the docking of KPNB1 to the NPC (Miorin *et al*., 2020). Conversely, Kato et al. suggested that aberrant nucleocytoplasmic transport is caused by the ORF6 dislocating NUP98 from the NPCs to the cytoplasm (Kato et al., 2021).

Full understanding of how ORF6 inhibits nucleocytoplasmic transport requires quantitative measurement of the dose response function, i.e., the relationship between the intracellular expression level of ORF6 and the resulting reduction in the nuclear transport efficiency. Previous studies measured neither of the quantities but instead assessed the average effect of ORF6 overexpression on the steady-state localization of IFN-related transcription factors or artificial cargoes or on the downstream IFN signaling activities. Accurate dose-response characterization would also enable objective comparison of ORF6 activity between viral species and measurement of perturbation effects, for example the influence of variations in ORF6 sequence on its inhibitory function. Such comparison may contribute to understanding of different characteristics between SARS-CoV-1 and SARS-CoV-2 and the prediction of the innate immune response to new SARS-CoV-2 variants which may occur in the future.

One reason that inhibition of nuclear transport by ORF6 was not accurately quantified in previous studies was the lack of suitable assays. We previously developed optogenetic assays for quantification of nuclear import and export in living cells (Yoo and Mitchison, 2021). Here, we apply them to measure inhibition by ORF6. We developed a simple measurement pipeline for calibrated quantification of the expression level of an untagged protein in single cells and applied it to measure ORF6 concentrations. In typical studies where results are averaged across many cells, variation in protein expression levels between cells is a technical problem. Conclusions based on bulk averages may not predict the behavior of individual cells whose expression level differs from the mean value. By measuring ORF6 expression in individual cells, we took advantage of this unavoidable variation to populate the concentration axis of dose-response plots. We applied the resulting dose-response technology to characterize and compare the inhibitory functions of artificial and natural variants of the ORF6 protein, revealing the difference between SARS-CoV-1 and SARS-CoV-2 ORF6s and providing new mechanistic insights for their inhibitory actions. Our live cell dose-response pipeline could be adapted to any measurement where expression of an exogenous protein alters some function that can be quantified in individual cells.

## Results

### Pipeline for live-cell dose-response characterization of nuclear transport inhibition

We sought to develop a method for measuring the dose-dependent effect of expressing a protein of interest (POI) on the nuclear transport kinetics in individual live cells, with the constrain that the POI is untagged. ORF6 is a small (MW ~7 kDa), membrane-associated protein, so tagging it with a fluorescent protein might perturb its activity. To measure nuclear transport kinetics, we adopted the optogenetics-based assays that we recently developed to measure the rates of classical nuclear import and export in live cells (Yoo and Mitchison, 2021). The assays use fluorescently labeled LOV2-based optogenetic probes that translocate from the cytoplasm to the nucleus or vice versa upon blue-light stimulation (Niopek et al., 2014; Niopek et al., 2016). The kinetics of the light-induced translocation is measured in each individual cell as proxy for the nuclear transport kinetics. The assays are performed using an automated microscopy platform and can readily process hundreds of cells in different conditions at different time points.

To express the POI and determine its level in individual cells while running the transport assay in parallel, we utilized a bicistronic expression system based on the 2A self-cleaving peptide (Szymczak and Vignali, 2005) (Figure 1A). U2OS cells stably expressing the optogenetic nuclear import or export probe were transfected with a plasmid encoding GFP-2A-POI. Due to the self-cleaving activity of the 2A sequence, which occurs during translation (de Felipe et al., 2006), GFP and POI are produced at the same rate as separate polypeptides. This enables the estimation of the POI concentration based on the GFP fluorescence intensity without the use of direct tagging which might disrupt the function of the POI. The GFP intensity and the transport kinetics were repeatedly measured in hundreds of individual cells at 3–6-hour intervals after transfection. The GFP intensity was converted to absolute GFP concentration via a novel calibration procedure, described below. The resulting data shows a wide range of GFP concentration, typically ranging from 0 to 30 μM when co-expressed with ORF6, due to the stochasticity in the transient transfection as well as the time-dependent increase in the expression level. Plotting the nuclear transport rate (import or export) against the GFP concentration revealed the dose-response curve. To generate parameters from these curves, we fit them to a Hill function. As in pharmacokinetic-pharmacodynamic modeling (Goutelle et al., 2008), this fit should be considered descriptive rather than mechanistic, but it allowed quantitative comparisons across all the data. Although GFP and POI are co-translated due to the 2A cleavage, their equilibrium concentrations may be different due to the difference in stability. When the absolute POI concentration is of interest, the GFP concentration can be converted into units of absolute protein concentration via a bulk biochemical analysis, for example quantitative western blot or mass spectrometry with spiked-in standards.

**Figure 1.**
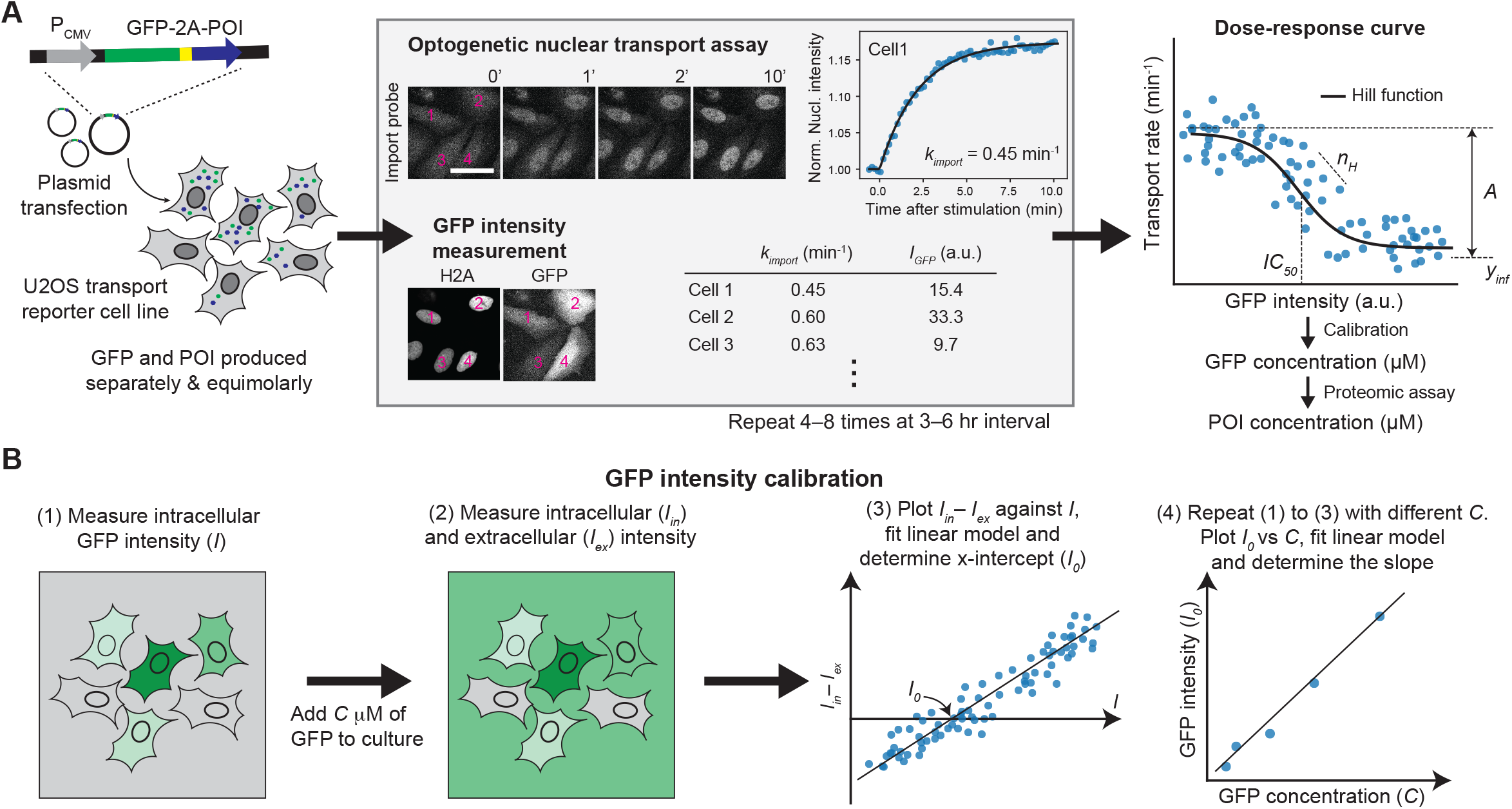
Calibrated dose-response characterization of nuclear transport alteration by exogenous protein expression. (A) U2OS cell line stably expressing H2A-Halo and optogenetic nuclear import (NES-mCherry-LINuS) or export probe (NLS-mCherry-NEXY) was transfected with a plasmid encoding GFP-2A-POI (protein of interest). Due to the 2A self-cleavage sequence, GFP and POI are produced at the same rate as separate polypeptides. The rate of the light-induced nuclear/cytoplasmic translocation of the transport probe and GFP intensity were simultaneously measured in individual cells. The measurement was repeated at 3–6-hr interval for ~24 hrs after the transfection. The GFP intensity was converted to GFP concentration via calibration. The Hill function was fitted to the relationship between the nuclear transport rate and GFP concentration (“dose-response curve”) to obtain the dose-response parameters. Scale bar 50 μm. (B) GFP intensity calibration procedure. (Step 1) Measure GFP intensity (*I*) of individual cells in the same way as in the dose-response characterization. (Step 2) Add a known concentration (*C* μM) of recombinant GFP to the culture and measure the intracellular and extracellular GFP intensities (*I*_*in*_ and *I*_*ex*_). (Step 3) Fit a linear model to the plot of *I*_*in*_ – *I*_*ex*_ vs *I* to determine the x-intercept (*I*_*0*_), which is the GFP intensity equivalent to the concentration *C*. (Step 4) Repeat Steps 1 to 3 with different *C*’s and fit a linear model to the plot of *I*_*0*_ vs *C* to determine the slope, which serves as the intensity-to-concentration conversion factor. See also Figure S1.

The GFP calibration was performed as follows (Figure 1B and S1): (Step 1) GFP expressing cells were imaged using the same microscope and image acquisition setting as in the dose-response characterization. The GFP intensity in each cell was quantified in the same way as in the dose-response characterization. (Step 2) Recombinant GFP was added to the culture at a certain concentration, and the same cells in the Step 1 were imaged again in the presence of GFP surrounding them. Cells could be brighter or darker than the background, depending on the relative concentration of the intracellular to the extracellular GFP concentration. (Step 3) Plotting the contrast between the intracellular and the background GFP intensity against the GFP intensity measured in Step 1 showed a linear relationship. A linear model was fitted to the relationship to determine the x-intercept, which corresponds to the GFP intensity of a cell whose intracellular GFP concentration is the same as the concentration of the recombinant GFP added to the culture in Step 2. (Step 4) Steps 1 to 3 were repeated with different concentrations of added GFP to obtain the relationship between the GFP intensity and concentration.

The slope of the linear model fitted to the relationship served as the intensity-to-concentration conversion factor. Importantly, this method reports GFP concentration values independent of the optical parameters of the microscope such as the depth of field or illumination brightness (Supplemental Text).

We tested the dose-response pipeline with negative controls (Figure S2). When only GFP was expressed, the nuclear import rate was constant across the GFP concentration, indicating the absence of artifacts due to GFP overexpression. ORF10 and Candidate ORF15 are gene products of SARS-CoV-2 that have been reported to not affect the IFN signaling and therefore are not likely to alter the nuclear import (Olson et al., 2021; Xia *et al*., 2020). As expected, these proteins showed flat dose-response curves, confirming that co-expression of GFP and an inactive POI does not perturb transport.

### Comparison of ORF6 activity between SARS coronaviruses

We applied the dose-response pipeline to compare the inhibitory effects of SARS-CoV-1 and SARS-CoV-2 ORF6s (denoted by ORF6^CoV1^ and ORF6^CoV2^, respectively) on nuclear transport. The amino acid sequences of ORF6^CoV1^ and ORF6^CoV2^ are 69% identical; most of the variations lie in the C-terminal half, including the 2 amino acid C-terminal extension of ORF6^CoV1^ (Figure 2A). We acquired dose-response curves of ORF6^CoV1^ and ORF6^CoV2^ on the nuclear import and export rates. The Hill function fit well to all the dose-response curves, providing parameters that reflect the amplitude of inhibition (*A*), half maximal inhibitory GFP concentration (*IC*_*50*_^GFP^), and the steepness of the dose-response curve (*n*_*H*_ or Hill coefficient) (Figure 2B, Table 1).

**Table 1.** Dose-response parameters of ORF6^CoV1^ and ORF6^CoV2^

| Construct | Transport direction | A (min<sup>-1</sup>) | IC<sub>50</sub><sup>GFP</sup> (μM) | IC<sub>50</sub><sup>ORF6</sup> (nM) | n<sub>H</sub> | y<sub>inf</sub> (min<sup>-1</sup>) |
| --- | --- | --- | --- | --- | --- | --- |
| GFP-2A-ORF6<sup>CoV1</sup> | Import | 0.43 ± 0.09 | 6.31 ± 2.18 | 94.7 ± 32.7 | 1.50 ± 0.45 | 0.16 ± 0.08 |
|  | Export | 1.34 ± 0.32 | 5.82 ± 2.40 | 87.3 ± 36.0 | 1.56 ± 0.62 | 1.13 ± 0.30 |
| GFP-2A-ORF6<sup>CoV2</sup> | Import | 0.47 ± 0.01 | 1.35 ± 0.07 | 6.08 ± 0.32 | 4.36 ± 0.77 | 0.15 ± 0.01 |
|  | Export | 1.43 ± 0.09 | 1.26 ± 0.18 | 5.67 ± 0.81 | 5.24 ± 3.39 | 1.01 ± 0.08 |

**Figure 2.**
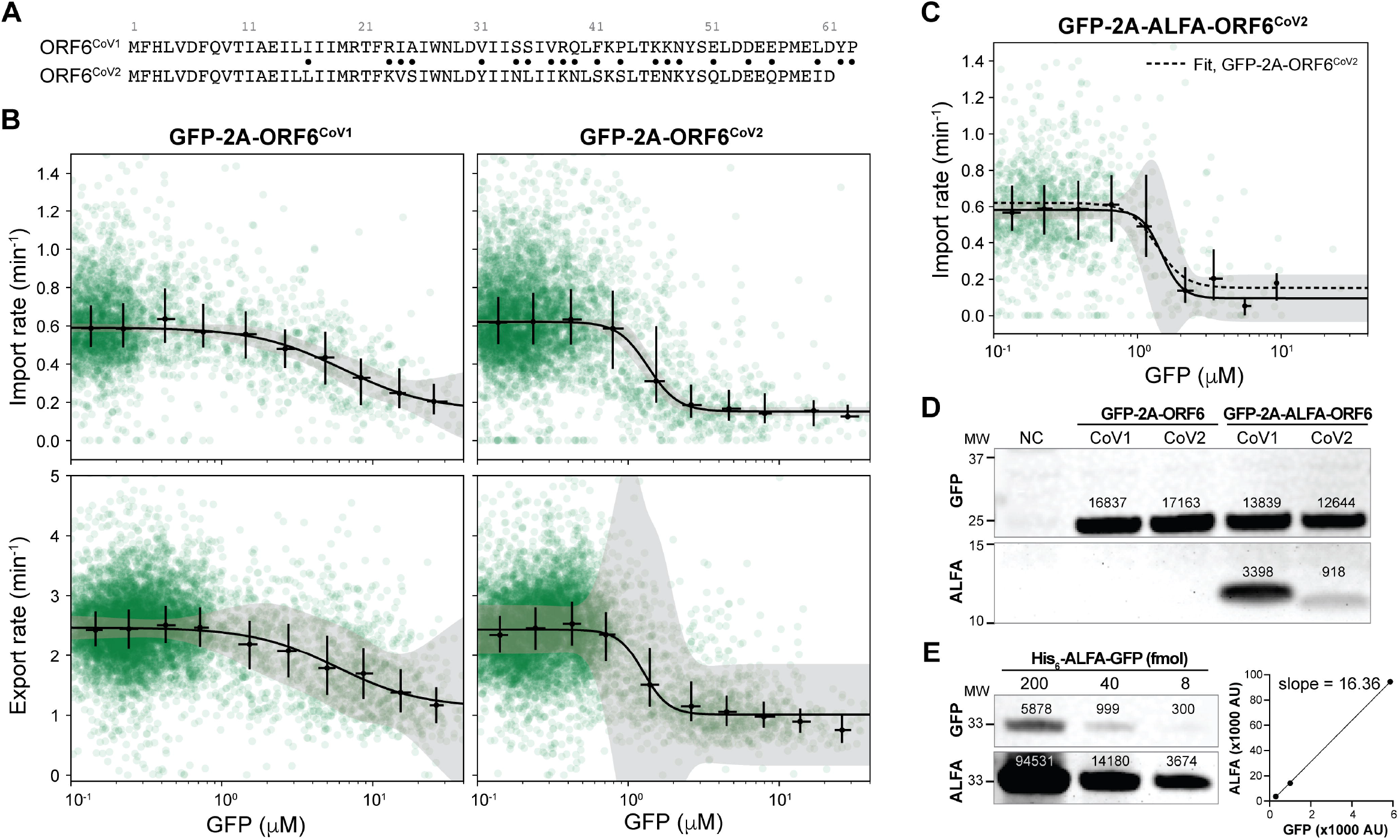
SARS-CoV-2 ORF6 is more potent than SARS-CoV-1 ORF6 in inhibiting bidirectional nuclear transport. (A) Amino acid sequences of SARS-CoV-1 and SARS-CoV-2 ORF6s (ORF6^CoV1^ and ORF6^CoV2^, respectively). Dots indicate the mismatches. (B) Dose-response curves for the nuclear import (top) and export (bottom) inhibition by ORF6^CoV1^ (left) and ORF6^CoV2^ (right). n>2900 cells for each combination. Green circles are the raw data points. Black error bar represents median and interquartile range in each bin. Black line and shaded area represent the best Hill function fit and corresponding 95% confidence interval, respectively. The parameter estimates and standard errors are shown in Table 1. See also Figure S3. (C) Dose response curves for the nuclear import inhibition by ALFA-ORF6^CoV2^ (n>1000 cells). The dashed line is the Hill function fit for the untagged ORF6^CoV2^ for comparison. See also Figure S4. (D) Anti-GFP (top) and anti-ALFA (bottom) western blot images of the lysate of U2OS cells transfected with nothing (NC), GFP-2A-ORF6s or GFP-2A-ALFA-ORF6s. Background subtracted integrated intensities shown above the bands. (E) Anti-GFP (left, top) and anti-ALFA (left, bottom) western blot images of the recombinant His_6_-ALFA-GFP with background subtracted integrated intensities shown above the bands. Linear regression of the anti-ALFA blot signal against the anti-GFP blot signal (right) results in the slope of 16.36. Combining the results in (D) and (E), we determined the molar ratio of GFP to ORF6 in the GFP-2A-ORF6 lysates to be 1000:15 for ORF6^CoV1^ and 1000:4.5 for ORF6^CoV2^.

In the nuclear import inhibition, ORF6^CoV1^ showed 4.7 times higher *IC*_*50*_^GFP^ than ORF6^CoV2^ (6.31 ± 2.18 μM vs 1.35 ± 0.07 μM) (Figure 2B, Table 1). The Hill coefficient was lower for ORF6^CoV1^ than ORF6^CoV2^ (1.50 ± 0.45 vs 4.36 ± 0.77), while the amplitude of inhibition was similar (0.43 ± 0.09 min^−1^ vs 0.47 ± 0.01 min^−1^). Both ORF6^CoV1^ and ORF6^CoV2^ inhibited the nuclear export, showing *IC*_*50*_ and Hill coefficient identical to those in the nuclear import inhibition (Figure 2B). This result indicates that ORF6 does not act specifically on the KPNB1-mediated classical nuclear import pathway but broadly affects nuclear import and export pathways mediated by multiple transport carriers. This is not consistent with the previously proposed model in which the nuclear transport inhibition depends on the specific interaction between ORF6 and KPNA2 (Frieman *et al*., 2007; Xia *et al*., 2020), while supporting models proposing NPC impairments (Addetia *et al*., 2021; Miorin *et al*., 2020). The nuclear import and export rates at each dose of ORF6^CoV1^ and ORF6^CoV2^ showed a colinear relationship, suggesting that these two different ORF6s use the same inhibitory mechanism (Figure S3).

The lower *IC* _*50*_ ^GFP^ of ORF6^CoV2^ could be due to a higher potency, a higher ORF6 concentration resulting from a higher protein stability, or the combination of both. Therefore, we sought to convert the *IC* _*50*_ ^GFP^ to the corresponding ORF6 concentration. Quantitative mass spectrometry was not successful due to the small size of the protein. We therefore used western blot with protein standards to measure the molar ratio of GFP and ORF6 co-expressed. Western blot using a commercial anti-ORF6 antibody was not successful in our hand (data not shown), so we tagged ORF6 with the ALFA-tag, a small epitope tag, and used the biotin-conjugated anti-ALFA nanobody for detection (Gotzke et al., 2019). N-terminus tagging of ORF6^CoV1^ and ORF6^CoV2^ with the ALFA-tag did not affect their dose responses of the nuclear import inhibition, suggesting that it does not perturb the inhibitory activity or the stability of ORF6 in cells (Figure 2C and S4). The lysate of U2OS cells transfected with GFP-2A-ORF6 or GFP-2A-ALFA-ORF6 showed a single band on the anti-GFP western blot at ~25 kD, indicating a high efficiency of the 2A self-cleavage (Figure 2D). Additional bands would have appeared at ~35 kD if the 2A self-cleavage had not been so efficient that the fusions of GFP and ORF6 had been produced. The anti-ALFA western blot of the lysate showed bands at ~10 kD region for the GFP-2A-ALFA-ORF6 transfected lysates but not for the GFP-2A-ORF6 transfected lysates, indicating that the anti-ALFA nanobody binding to ORF6 is negligible (Figure 2D). The ratios of the anti-GFP to anti-ALFA blot intensities were 0.246 for ALFA-ORF6^CoV1^ and 0.073 for ALFA-ORF6^CoV2^, suggesting a significantly lower stability of ORF6^CoV2^. On the same anti-GFP and anti-ALFA western blots, recombinant His_6_-ALFA-GFP appeared at ~33 kD with the intensity ratio of 1 to 16.36 (Figure 2E). Using this to correct for the difference in the intensities between anti-GFP and anti-ALFA blots against an equal molar amount of antigens, we determined the molar ratio of GFP and ALFA-ORF6 to be 1000 to 15 for ORF6^CoV1^ and 1000 to 4.5 for ORF6^CoV2^. Consequently, we estimated the half maximal inhibitory ORF6 concentrations (*IC*_*50*ORF6_) to be ~90 nM for ORF6^CoV1^ and ~6 nM for ORF6^CoV2^ (Table 1), indicating that the inhibitory potency of ORF6^CoV2^ is ~15 times higher. The higher potency of ORF6^CoV2^ in the nuclear import inhibition may underlie the previous observation that ORF6^CoV2^ suppressed IFN signaling more strongly than ORF6^CoV1^ when a cell population was transfected with the same amount of ORF6-expressing plasmid (Kimura *et al*., 2021; Olson *et al*., 2021).

We used enhanced GFP (EGFP) for the dose-response measurements throughout this study but also tested two other GFP variants with different maturation times and spectral characteristics, mCerulean and mTurquoise2 (Balleza et al., 2018), and obtained very similar dose-response curves for ORF6^CoV2^ (Figure S5). This suggests that the dose-response pipeline is not sensitive to the choice of the fluorescent protein, further supporting its broad applicability.

### The absence of C-terminal Tyr-Pro extension largely explains the higher potency of SARS-CoV-2 ORF6

ORF6 is characterized by abundance of hydrophobic residues in the N-terminal region and negatively charged residues in the C-terminal region (Figure 3A). To identify regions in ORF6 that are accountable for the difference in the dose-response characteristics between ORF6^CoV1^ and ORF6^CoV2^, we performed site-directed mutagenesis to substitute each cluster of mismatches in ORF6^CoV2^ with corresponding ORF6^CoV1^ residues (Figure 3A). None of the mutations affected the amplitude of inhibition (*A*), while some of them significantly altered the *IC*_*50*_ ^GFP^ and Hill coefficient (*n* _*H*_) of ORF6 (Figure 3B and S6).

**Figure 3.**
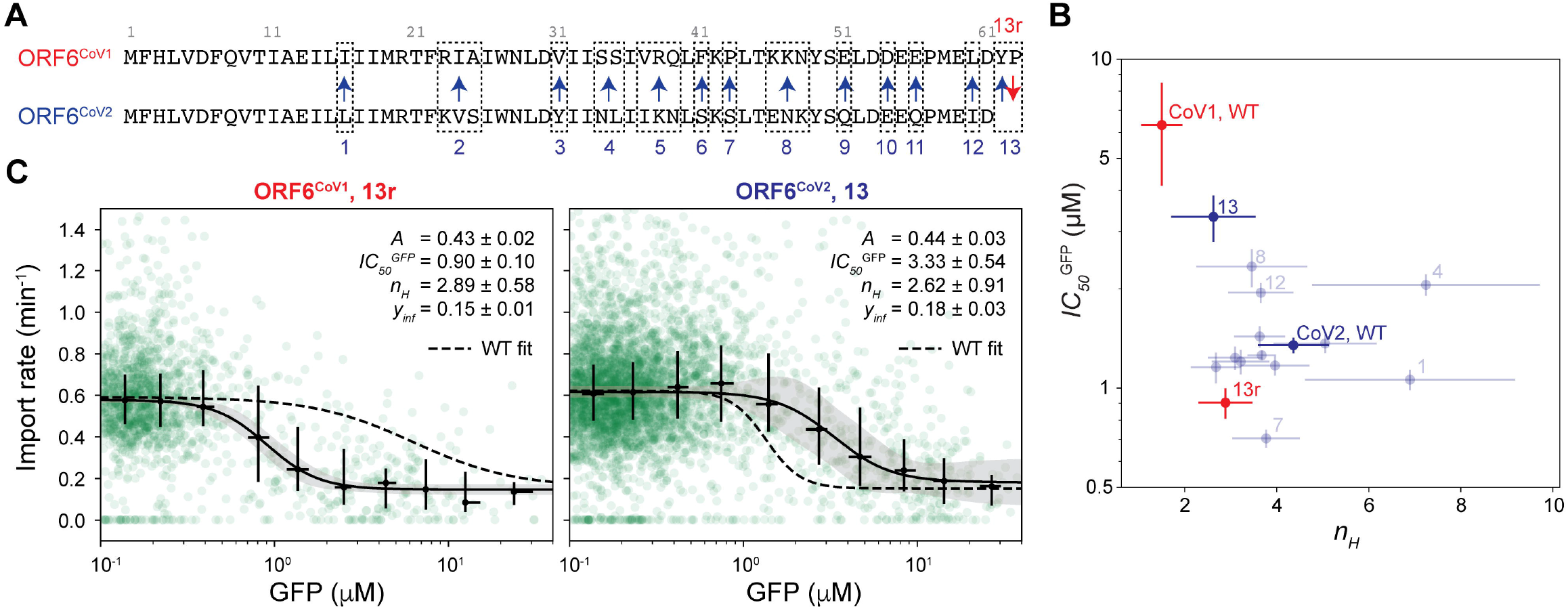
The lack of C-terminal Tyr-Pro mainly contributes to the higher potency of SARS-CoV-2 ORF6. (A) Notation of ORF6^CoV1^ and ORF6^CoV2^ variants. (B) Plot of *IC* _*50*_ ^GFP^ vs *n* _*H*_ for each ORF6 variant. Error bar represents the standard error of the parameter estimate. n>1000 cells for each variant. (C) Dose-response curves for the nuclear import inhibition by #13r (deletion of YP from the C-terminal end of ORF6^CoV1^, left) and #13 (addition of YP to the C-terminus of ORF6^CoV2^, right). Green circles are the raw data points. Black error bar represents median and interquartile range in each bin. Black line and shaded area represent the best Hill function fit and the corresponding 95% confidence interval, respectively. The parameter estimates and standard errors are shown in the plots. Dotted line is the best Hill function fit for the ORF6^CoV1^ (left) and ORF6^CoV2^ (right) wild types shown for comparison. See also Figure S6.

Notably, three mutations, E46K/N47K/K48N (#8), I60L (#12) and C-terminal addition of YP (#13), changed both the *IC* _*50*_^GFP^ and *n*_*H*_ of the ORF6^CoV2^ toward those of ORF6^CoV1^ (Figure 3B and S6). Among them, the mutation #13 showed the strongest effect, increasing *IC*_*50*_^GFP^ by a factor of 2.5 and decreasing Hill coefficient by a factor of 1.7 (Figure 3C). We verified the effect of this mutation by removing YP at the C-terminus of ORF6^CoV1^ (#13r). The mutation #13r drastically reduced the *IC*_*50*_^GFP^ of ORF6^CoV1^ by a factor of 7, making it even lower than ORF6^CoV2^ (Figure 3B and 3C). This suggests that the lack of C-terminal YP mainly contributes to the higher potency of ORF6^CoV2^. Consistent with our results, a previous study observed that mutations #8 and #13 significantly decreased the inhibitory effect of ORF6^CoV2^ on the IFN signaling activity (Kimura *et al*., 2021).

### The N-terminal and C-terminal regions are simultaneously required for the inhibitory function of SARS-CoV-2 ORF6

To gain more insights into the inhibitory mechanism, we characterized additional artificial and natural variants of ORF6^CoV2^. The hydrophobic N-terminal region of ORF6 interacts with the membranes of ER and Golgi apparatus; it has been previously shown that residues 2-37 of ORF6^CoV1^ are embedded in the membrane (Zhou et al., 2010) and that the peptides of residues 4-24 and 22-42 of ORF6^CoV2^ bind the membrane (O’Keefe et al., 2021). The negatively charged C-terminal region is responsible for interactions with NUP98-RAE1 and KPNAs (Addetia *et al*., 2021; Frieman *et al*., 2007; Kimura *et al*., 2021; Miorin *et al*., 2020). Therefore, it is predicted that mutations in the C-terminal region of ORF6 would affect its interactions with NUP98-RAE1 and/or KPNA while retaining the membrane binding.

We found that C-terminal truncation of 24 residues resulted in the complete loss of the inhibitory effect of ORF6^CoV2^ on the nuclear import, and so did C-terminal 6-residue truncation, which had been discovered in COVID-19 patients in Italy (Delbue et al., 2021) (Figure 4A). Together with the comparative mutational analysis of ORF6^CoV1^ and ORF6^CoV2^ (Figure 3), these results suggest that the C-terminal end is crucial to the inhibitory function of ORF6. We also tested a point mutation, M58R, which has been shown to abolish the ability of ORF6^CoV2^ to bind NUP98-RAE1 while maintaining its KPNA binding (Miorin *et al*., 2020). This point mutation also resulted in the complete loss of the inhibitory function of ORF6^CoV2^, suggesting that NUP98-RAE1 binding of the C-terminal region is essential while KPNA binding is not (Figure 4A).

**Figure 4.**
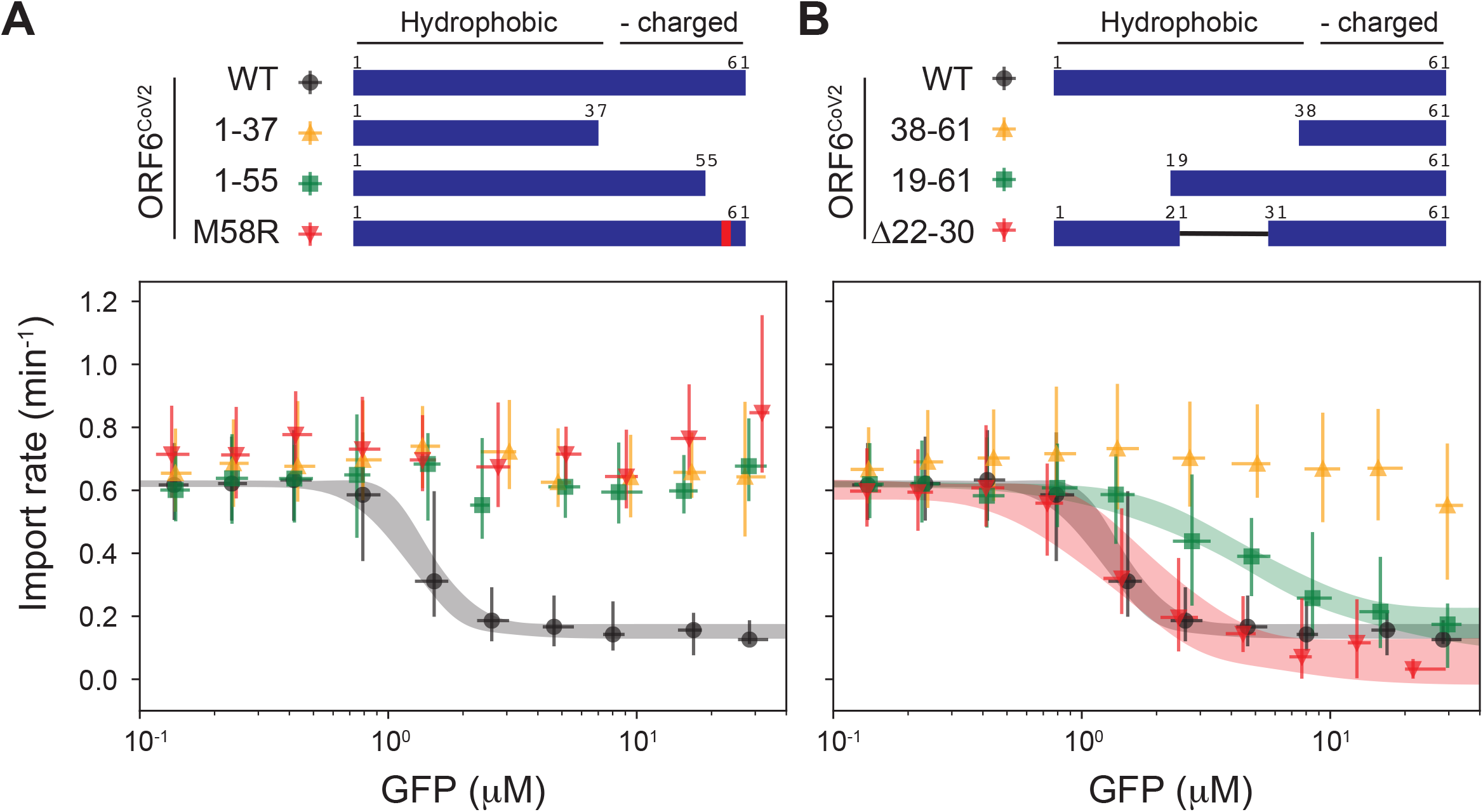
The N-terminal and C-terminal regions are simultaneously required for ORF6 activity. (A and B) Dose-response curves for the nuclear import inhibition by (A) ORF6^CoV2^ WT (black circles), residues 1-37 (yellow upward-pointing triangles), residues 1-55 (green squares), and M58R mutant (red downward-pointing triangles); and (B) WT (black circles), residues 38-61 (yellow upward-pointing triangles), residues 19-61 (green squares), and Δ22-30 mutant (red downward-pointing triangles). n>1100 cells for each variant. Error bars show the median and interquartile range in each bin. Shaded area represents 95% confidence interval of the best Hill function fit. Fits are not shown for the dose-response curves that failed to reject the null hypothesis that the inhibition amplitude (*A*) is zero (α= 0.05).

We also evaluated the influence of mutations in the N-terminal and middle regions on the inhibitory function of ORF6^CoV2^ (Figure 4B). The peptide of C-terminal residues 43-61 has been previously shown to interact with NUP98-RAE1 complex (Miorin *et al*., 2020). We found that the peptide of residues 38-61 did not show a significant inhibitory effect on the nuclear import (Figure 4B), indicating that NUP98-RAE1 binding alone is not sufficient for the transport inhibition. Alternative translation of ORF6^CoV2^ gene results in the lack of the first 18 residues (Finkel et al., 2021), which we found to increase the *IC* _*50*_ ^GFP^ of ORF6^CoV2^ by a factor of 3.4 (4.57 ± 0.76 μM) (Figure 4B). A previous study discovered ORF6 having a 9-residue deletion from the central part (residues 22-30) in a SARS-CoV-2 strain passaged *in vitro* in the IFN-deficient Vero E6 cell (Riojas et al., 2020). Although the authors predicted that the deletion would dramatically alter the structure of ORF6 binding to the membrane, the deletion did not affect the dose-response characteristics of the nuclear import inhibition (Figure 4B). This is consistent with K23R/V24I/S25A and Y31V mutations (denoted by #2 and #3) not affecting the dose-response characteristics (Figure 3 and S6). Taken together, these mutational analyses suggest that the N-terminal and C-terminal regions are simultaneously required for the ORF6 activity in the nuclear transport inhibition, while the central region (22-30) is not important.

### Membrane binding is not required for the inhibitory function of ORF6

We next sought to examine the localization of ORF6 using small epitope tags and immunofluorescence. Direct tagging of ORF6 with an epitope tag or fluorescent protein has been performed in previous studies to examine the subcellular localization or interactions with other proteins (Addetia *et al*., 2021; Frieman *et al*., 2007; Gordon *et al*., 2020; Kato *et al*., 2021; Kimura *et al*., 2021; Miorin *et al*., 2020; Xia *et al*., 2020), but the effect of the tagging on the inhibitory function has not been evaluated. We characterized the dose responses of ORF6^CoV2^ N- or C-terminally tagged with three different epitope tags: ALFA-tag (Gotzke *et al*., 2019), Flag-tag, and HA-tag. All three epitope tags increased the *IC* _*50*_ ^GFP^ of ORF6^CoV2^ when positioned at the C-terminus (Figure S7). This is consistent with the observation that the *IC* _*50*_ ^GFP^ of ORF6^CoV2^ was sensitive to the C-terminal modifications (Figure 3 and 4A). On the other hand, at the N-terminus of ORF6, ALFA-tag and HA-tag had negligible influence on the dose-response characteristics, while Flag-tag rather reduced the *IC*_*50*_^GFP^ (Figure S7). These results suggest that the tagging location and the type of epitope tag should be carefully selected when studying small proteins and must be explicitly reported in publications.

We chose to use ALFA-tag for subsequent immunofluorescence analyses since there is a well-characterized fluorophore-conjugated anti-ALFA nanobody commercially available (Gotzke *et al*., 2019). U2OS cells were transfected with GFP-2A-ALFA-ORF6^CoV2^ or GFP-2A-ORF6^CoV2^-ALFA for 24 hours. Prior to paraformaldehyde fixation, GFP concentration was measured to estimate the co-expressed ORF6 level as in the nuclear transport dose-response analysis (Figure 1B). The fixed cells were permeabilized with digitonin to preserve the intracellular membrane structure and then immunostained using the anti-ALFA nanobody and anti-NUP98 antibody. Interestingly, the N- and C-terminally tagged ORF6^CoV2^ showed distinct staining patterns. ALFA-ORF6^CoV2^ showed strong nuclear pore staining with negligible staining elsewhere throughout different expression levels (Figure 5A). On the other hand, ORF6^CoV2^-ALFA was stained at the cytoplasmic membranes and the nuclear envelope so strongly that staining of individual nuclear pores was barely discernable (Figure 5B). At high ORF6 concentrations, we found that NUP98 was dislocated from the NPC, consistent with a previous study (Kato *et al*., 2021). However, the NUP98 dislocation was not noticeable at ORF6 concentrations that are low yet high enough to inhibit the nuclear transport, so it is unlikely to be directly involved in the nuclear transport inhibition.

**Figure 5.**
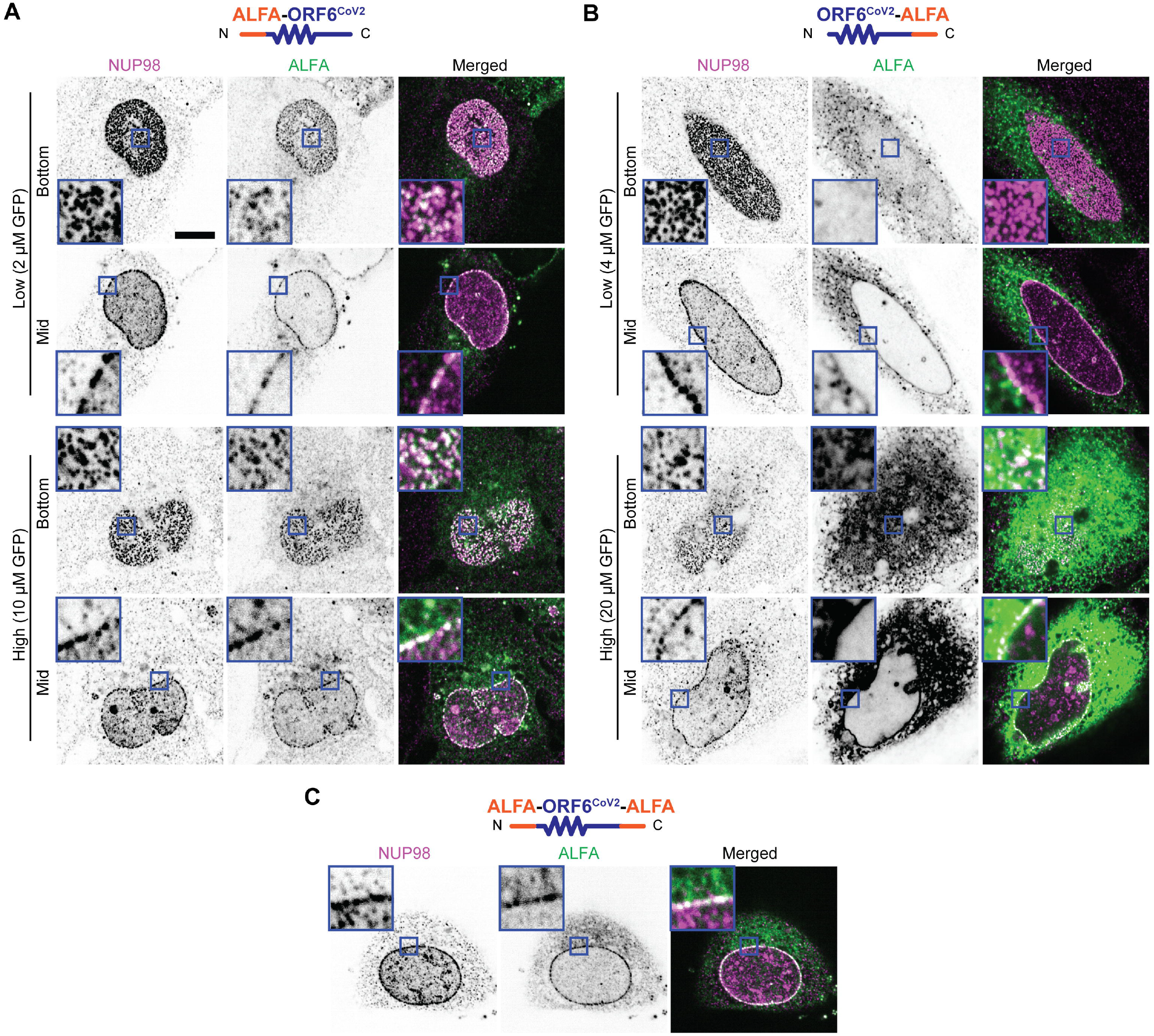
Membrane binding is not required for the inhibitory function of ORF6. (A, B) U2OS cells were transfected with GFP-2A-ALFA-ORF6^CoV2^ or GFP-2A-ORF6^CoV2^-ALFA so that ORF6^CoV2^ N- or C-terminally tagged with ALFA was co-expressed with GFP. Representative NUP98 and ALFA immunofluorescence images of (A) ALFA-ORF6^CoV2^ and (B) ORF6^CoV2^-ALFA at low and high GFP concentrations when focused on the basal plane (“bottom”) and midplane (“mid”) of the nucleus. See also Figure S7. (C) Representative NUP98 and ALFA immunofluorescence image of ALFA-ORF6^CoV2^-ALFA when focused on the midplane of the nucleus. Scale bars 10 μm. Insets: 10x magnification.

The difference in the ALFA immunostaining patterns could result from the membrane binding of the N-terminal region being disrupted by the proximal N-terminal ALFA-tag but not by the distal C-terminal tag. If this was the case, the lack of effect of the N-terminal ALFA-tag on the dose-response characteristics (Figure S7) would suggest that membrane binding is not required for the inhibitory function of ORF6^CoV2^. However, the difference in the ALFA immunostaining patterns could also result from the membrane binding of the N-terminal region restricting the N-terminal ALFA tag from binding the anti-ALFA nanobody. To test this possibility, we examined the immunofluorescence localization of ORF6 having ALFA tags at both the N- and C-termini (ALFA-ORF6^CoV2^-ALFA) (Figure 5C). We found that this construct showed strong ALFA immunostaining at the nuclear pores, similar to ALFA-ORF6^CoV2^. If the absence of the immunostained ALFA-ORF6^CoV2^ at the membranes was due to epitope inaccessibility rather than hindrance of membrane binding, ALFA-ORF6^CoV2^-ALFA would have shown strong ALFA immunostaining at the membranes like ORF6^CoV2^-ALFA did. Thus, the N-terminal ALFA tag inhibits the membrane binding of ORF6^CoV2^ but does not alter its inhibitory effect on the nuclear transport. This indicates that the role of the N-terminal region in the nuclear transport inhibition is not the membrane binding but something else.

### Forced homo-oligomerization rescues the inhibitory function of N-terminally truncated ORF6

We next sought to investigate alternative roles of the N-terminal region of ORF6 in the transport inhibition. Several other proteins have also been reported to bind FG-NUPs and inhibit the carrier-mediated nuclear transport. Such proteins include wheat germ agglutinin (Finlay et al., 1987) (WGA), hyperphosphorylated tau (Eftekharzadeh et al., 2018), C9orf72 dipeptide repeats (Shi et al., 2017), and vesicular stomatitis virus matrix (VSV M) protein (Faria et al., 2005; Petersen et al., 2000). A common feature of these proteins is the ability to form protein aggregates or homo-oligomers which are presumably able to multivalently cross-link FG domains (Gaudin et al., 1995; Gotz et al., 2019; Mori et al., 2013). For example, WGA forms a dimer with eight binding sites for O-GlcNAc, a glycosylation abundant in the FG domains (Li and Kohler, 2014; Schwefel et al., 2010). A recent computational analysis predicted that ORF6 is also aggregate-prone due to the hydrophobic N-terminal region (Flores-Leon et al., 2021). Therefore, we hypothesized that the mechanistic role of the N-terminal region of ORF6 in the nuclear transport inhibition is to mediate aggregation or homo-oligomerization, while its membrane binding is rather coincidental.

To test this hypothesis, we investigated whether the N-terminal region can be functionally replaced with an oligomerization domain derived from other proteins that do not bind membranes. We first used a chemically inducible oligomerization system based on the F36V mutant of FK506-binding protein 12 (FKBP) and the B/B Homodimerizer ligand (equivalent to AP20187) (Clackson et al., 1998) (Figure 6A). Addition of the homodimerizer did not affect the dose-response characteristics of the full-length ORF6^CoV2^ or the N-terminal truncation of ORF6^CoV2^ (residues 38-61; CT), confirming the absence of unspecific effects of the homodimerizer on the dose response (Figure 6A). When the CT was fused to FKBP, addition of the homodimerizer partially recovered the inhibitory function of the CT (Figure 6A).

**Figure 6.**
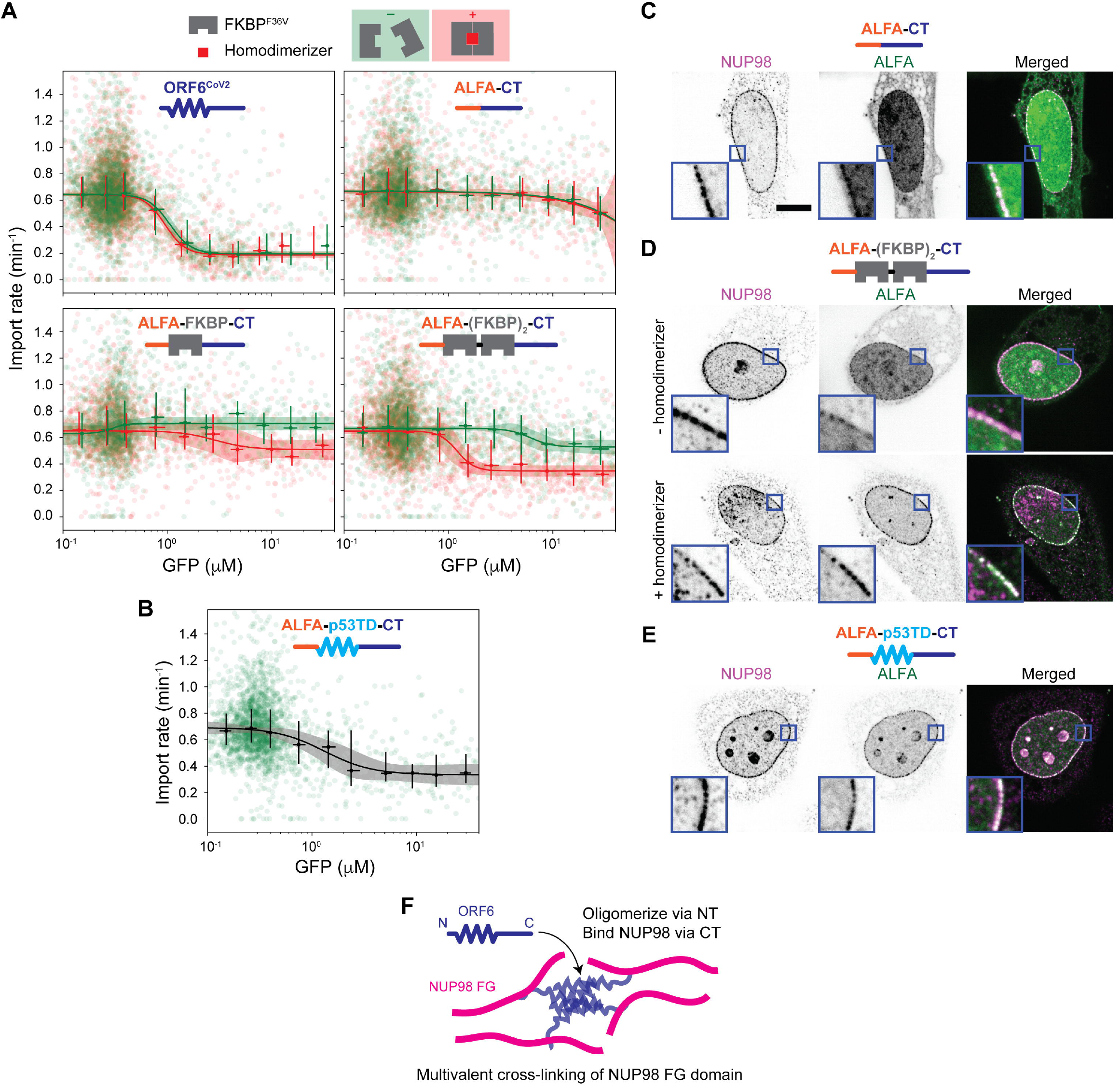
Forced homo-oligomerization rescues the inhibitory function of the N-terminally truncated ORF6. (A) FKBP homo-dimerizes in the presence of the B/B homodimerizer. Dose-response curves for the full-length ORF6^CoV2^ (top left), ALFA-CT (top right), ALFA-FKBP-CT (bottom left), and ALFA- (FKBP)_2_-CT (bottom right), where the CT refers to the C-terminal region (residue 38-61) of ORF6^CoV2^. Green and red colors correspond to the absence and presence of 500 nM B/B homodimerizer, respectively. (B) The dose-response curve for ALFA-p53TD-CT, where p53TD is the tetramerization domain (residues 326-356) of p53. In A and B, n>1500 cells for each condition. Error bar shows median and interquartile range in each bin, and the solid line and shaded area represent the best Hill function fit and the corresponding 95% confidence interval, respectively. (C, D, and E) NUP98 and ALFA immunofluorescence images of cells transfected with (C) ALFA-CT, (D) ALFA-(FKBP)_2_-CT in the absence (top) and presence (bottom) of 500 nM B/B homodimerizer, and (E) ALFA-p53TD-CT. Samples were focused on the midplane of the nucleus. Scale bar 10 μm. Insets: 10x magnification. (F) Proposed model. The N-terminal region (NT) of ORF6 drives the oligomerization, leading to cross-linking of the FG domains of NUP98 at the nuclear pores via multivalent binding of the C-terminal region (CT).

The homodimerizer-dependent recovery further increased when the CT was fused to two FKBPs in tandem, which presumably induce higher-order oligomerization (Figure 6A). We also tested the tetramerization domain of p53 (residues 326-356; p53TD) (Clore et al., 1994; Jeffrey et al., 1995), which is far smaller than the tandem FKBP (MW ~4 kDa vs ~24 kDa) and comparable to the truncated N-terminal region in size. We found that fusing p53TD also rescues the inhibitory function of the CT (Figure 6B). We confirmed that the CT and the fusion proteins were localized at the nuclear pores without showing noticeable membrane localization (Figure 6C, 6D, and 6E). The recovery of the inhibitory function via forced homo-oligomerization suggests that the mechanistic role of the N-terminal region in the nuclear transport inhibition may be to drive oligomerization of ORF6 and that membrane binding is dispensable (Figure 6F).

Having multiple NUP98-binding C-terminal regions, the ORF6 oligomer would have a high avidity for the multiple copies of NUP98 clustered at the NPC. The interaction with KPNAs, on the other hand, would not be strengthened by homo-oligomerization because they diffuse freely as monomers. Consistent with this prediction, we found that forced homo-oligomerization increased the nuclear pore localization of the CT relative to the nucleoplasmic localization, which presumably results from the KPNA binding (Figure 6C, 6D, and 6E). Thus, we propose that ORF6 homo-oligomerizes to multivalently cross-link the FG domains of NUP98 at the NPC (Figure 6F).

## Discussion

We developed a broadly applicable pipeline for live-cell dose-response characterization of nuclear transport inhibitions. This combines the optogenetics-based measurement of nuclear transport kinetics (Yoo and Mitchison, 2021) with measurement of protein expression level in individual cells. The protein level measurement does not rely on direct tagging and therefore is suitable for small proteins like SARS coronavirus ORF6. Both dose and response are calibrated in physical units, μM and min^−1^, making the data reproducible between experiments and directly comparable across different conditions and microscopes. Here we chose to fit the widely used Hill function to the dose-response data to obtain quantitative metrics, noting that more mechanistic models may also be used to extract biophysical parameters in future studies.

The dose-response pipeline quantitatively revealed key mechanistic features of the inhibitory function of ORF6. First, ORF6 inhibits multiple carrier-mediated bidirectional transport pathways. ORF6 showed the same potency and Hill coefficient for inhibitions of KPNB1-mediated nuclear import and CRM1/XPO1-mediated nuclear export (Figure 2B). Moreover, a recent study showed that ORF6 disrupts the nuclear export of mRNA (Addetia *et al*., 2021), which is mediated by a variety of other export carriers (Carmody and Wente, 2009). Therefore, the nuclear transport inhibition likely arises from a carrier unspecific perturbation of the nuclear transport machinery, e.g., the NPC impairment, rather than from specific interactions with KPNAs. Second, NUP98 binding of the C-terminal region is necessary but not sufficient for the inhibitory action of ORF6. C-terminal modifications dramatically altered the dose-response characteristics of ORF6 (Figure 3, 4 and 5). Most importantly, the NUP98 binding deficient mutation, M58R (Miorin *et al*., 2020), abolished the inhibitory function of ORF6 (Figure 4A). Compared to ORF6^CoV1^, ORF6^CoV2^ showed a higher potency in inhibiting the nuclear transport in our study (Figure 2B), primarily due to differences in the C-terminal end. In another study, ORF6^CoV2^ showed a stronger NUP98 binding than ORF6^CoV1^ (Addetia *et al*., 2021). This correlation suggests that NUP98 binding of the C-terminal region determines the potency of ORF6. The NUP98 binding C-terminal region alone without the N-terminal region was not able to inhibit the nuclear transport, indicating that the ORF6 function requires the joint action of N- and C-terminal regions (Figure 5). Finally, membrane binding of the N-terminal region is not required for the inhibitory function of ORF6. Disrupting the membrane binding by N-terminally tagging the ORF6^CoV2^ with ALFA did not alter the inhibitory dose-response characteristics (Figure 5). The N-terminal region was able to be functionally replaced with oligomerization domains that do not bind membranes (Figure 6).

Therefore, we argue that the major role of the N-terminal region in the nuclear transport inhibition is to drive oligomerization, while the membrane binding is dispensable. Taken together, we propose that ORF6 forms oligomer via the N-terminal region to multivalently cross-link the FG domains of NUP98 via the C-terminal region (Figure 6F). The long FG domain of NUP98 was found to interact *in vivo* with many other FG-NUPs of native human NPCs (Ma et al., 2016), so the cross-linking of the FG domains of NUP98 could have strong influence on the overall permeability of the FG barrier to cargo-carrier complexes.

Blocking nuclear transport is an interesting strategy for RNA viruses to evade innate immunity. If this block is too rapid or too potent, viral replication might be compromised by depletion of host cytoplasmic factors needed for translation and replication. SARS-CoV-1 and SARS-CoV-2 appear to have evolved to different optima, where the blocking function of ORF6 is much stronger in SARS-CoV-2. This presumably increases the ability of SARS-CoV-2 to evade interferon signaling, possibly at some cost in replication efficiency that might be compensated by other adaptations. How these differences contribute to the overall pathogenicity of SARS-related viruses is an important topic for future research.

## Acknowledgments

We thank Prof. Pamela Silver (Harvard Medical School, Boston, MA) and the members of the Silver laboratory, especially Rui Tong Quek and Dr. Tai Ng, for providing plasmids, reagents, and comments; Dr. James Pelletier (Centro Nacional de Biotecnología, Madrid, Spain) for sharing the idea on GFP intensity calibration; and Dr. Luke Lavis (Janelia Research Campus, Ashburn, VA) for providing JF646 dye. We also thank the Nikon Imaging Center (Harvard Medical School) for help with light microscopy.

This study was supported by NIH grant R35GM131753, a sponsored research award from AbbVie Inc. and postdoctoral fellowship (T.Y.Y) F32GM131585.

## Author contributions

T.Y.Y. and T.J.M. conceived the study and wrote the manuscript. T.Y.Y. designed and performed the experiments and analyzed the data with input from T.J.M.

## Methods

### Cell culture

U2OS cell lines (engineered from HTB-96, ATCC) were maintained in complete DMEM (low glucose (1g/L) Dulbecco’s modified Eagle’s medium (DMEM, Thermo Fisher, #10567022) supplemented with 10% Fetal Bovine Serum (FBS, Thermo Fisher, #A31605), 50 IU ml^−1^ penicillin, and 50 μg ml^−1^ streptomycin (Thermo Fisher, #15140122)) at 37°C in a humidified atmosphere with 5% CO_2_. Cells were validated as mycoplasma free by PCR-based mycoplasma detection kit (ATCC, #30-1012K).

### Plasmid cloning and reagent preparation

Plasmids encoding GFP-2A-ORF6 were generated by inserting EGFP-T2A sequence (copied from Addgene #140424 by PCR) at the 5’ end of the ORF6 gene in pSecTag2 mammalian expression vector using Gibson Assembly. ORF6 mutants were generated by using Q5 Site-Directed Mutagenesis Kit (NEB, #E0554S) or QuikChange II XL Site-Directed Mutagenesis Kit (Agilent, #200521). ALFA-tag (PSRLEEELRRRLTEP), HA-tag and Flag-tag were attached to the N- and/or C-terminal end of ORF6 with Gly-Ser linker. Oligonucleotides for PCRs were purchased from IDT or Genewiz. All the plasmids generated in this study were verified by Sanger sequencing (Genewiz).

Recombinant His_6_-EGFP used in the GFP intensity calibration (Figure 1B) was expressed in Rosetta 2 (DE3) competent cells (Millipore Sigma #71400) using pDual-EGFP plasmid (Addgene, #63215), purified using HisPur™ Ni-NTA Spin Columns (Thermo Fisher, #88226) via a standard protocol, and dialyzed in 50 mM Tris-HCl pH 7.5, 150 mM NaCl, 10% glycerol, 2 mM DTT. The concentration was determined based on the 488 nm absorbance and the previously reported value (Shimizu et al., 2017) of the extinction coefficient of EGFP at 488 nm, which is 53,300 M^−1^cm^−1^. The purified His_6_-EGFP was aliquoted, snap-frozen in liquid nitrogen, and stored in −80°C. Recombinant His_6_-mCerulean and His_6_- mTurquoise2 were prepared similarly using pBAD-mCerulean and pBAD-mTurquoise plasmids (Addgene, #54666 and #54844) and TOP10 competent cells (Thermo Fisher, #C404010).

### Dose-response characterization of nuclear transport inhibition

#### Live-cell imaging

U2OS stable cell lines stably expressing H2A-Halo and NES-mCherry-LINuS (import probe) or NLS-mCherry-LEXY (export probe) were generated in our previous study (Yoo and Mitchison, 2021) and were used in this study for measuring the dose-dependent effect of ORF6 on the nuclear transport kinetics. Cells were seeded at ~10,000 cells per well in an eight-well chambered coverslip (ibidi, #80826) with 250 μl complete DMEM. On the next day, cells in each well were transfected with a plasmid encoding GFP-2A-POI (protein of interest) using TransIT-2020 transfection reagent (Mirus #5404) as follows: 260 ng plasmid was complexed with 0.78 μl reagent in 26 μl Opti-MEM™ I Reduced Serum Medium (Thermo Fisher, #31985062) for 20 minutes. The complex was diluted in 263 μl prewarmed imaging media (low glucose (1g/L) DMEM without phenol red (Thermo Fisher, #11054020) supplemented with 10% FBS, 50 IU ml^−1^ penicillin, 50 μg ml^−1^ streptomycin, and GlutaMAX Supplement (Thermo Fisher, #35050061)) containing 500 nM JF646-HaloTag ligand (gift from Luke Lavis) and added to the well. For the FKBP-based forced oligomerization experiments (Figure 6A), 500 nM B/B Homodimerizer (Takara, #635058) was also added. After 2.5–6 hours of the transfection, cells were continually imaged for the simultaneous measurement of the nuclear transport kinetics and GFP intensity for ~24 hours. Cage microscope incubator (OkoLab) was used to maintain the cells at 37°C in 5% CO_2_ with high humidity during imaging.

Live-cell imaging for the simultaneous measurement of the nuclear transport kinetics and GFP intensity was performed on a Nikon Ti motorized inverted microscope with Perfect Focus System, using spinning disk confocal scanner (CSU-X1, Yokagawa) with Spectral Applied Research Aurora Borealis modification, motorized stage and shutters (Proscan II, Prior), scientific complementary metal-oxide-semiconductor (sCMOS) camera (Flash4.0 V3, Hamamatsu), laser merge module (LMM-5, Spectral Applied Research), CFI Plan Apo 20x/0.75NA objective lens (Nikon), and ZT445/514/561/640tpc (Chroma) polychroic mirror. mCherry-labeled import/export probes were imaged using 561-nm laser and ET605/70m emission filter (Chroma), while H2A-Halo:JF646 was imaged using 642-nm laser and ET700/75m emission filter (Chroma). Live-cell time-lapse images were acquired through a ~11-min imaging cycle that consisted of two acquisition phases: pre-activation and activation. Throughout the cycle, the mCherry-labeled transport probes and H2A-Halo:JF646 were imaged every 10 s with 100-ms exposure time. Three frames (30 s) were acquired in the pre-activation phase to determine the baseline nuclear localization of the probe, followed by a 10-min activation phase (61 frames) in which the optogenetic transport probes were activated by 200-ms exposure to the activation laser (447 nm) every 10 s. The activation laser also excites GFP, of which images were collected using ET480/40m emission filter (Chroma). To increase the throughput, the imaging cycle was executed at three different fields simultaneously. The imaging cycle was repeated at 3–6-hour interval at different wells, which was fully automated by using Journal macro in MetaMorph Software. The power of the 447-nm activation laser was optimized based on the nuclear export rate vs. laser power curve, to the lowest saturation level where a small variation of the power does not influence the transport rate. No sign of photobleaching or photodamaging was noticed when the imaging cycle was repeated every hour for 24 hours at the same field.

#### Computational image analysis

Custom Python codes were used for quantitative image analysis of the time-lapse images. The quantification of the nuclear transport kinetics was performed as described previously (Yoo and Mitchison, 2021). Briefly, cell nuclei were segmented based on H2A-Halo:JF646 images using U-Net convolutional neural network trained on manually annotated U2OS nuclei images (Caicedo et al., 2019) and tracked using Trackpy package (v0.4.1, 10.5281/zenodo.1226458). Then, for each nucleus, the time-trajectory of the mean nuclear mCherry intensity was measured and normalized such that the average intensity during pre-activation phase was 1 and the background intensity was 0. The following monoexponential decay model was fitted to the normalized nuclear mCherry intensity trajectory in the activation phase to determine the nuclear transport rate *k*:

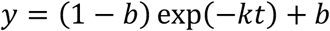

When the import probe is used, *k* corresponds to the import rate and *b* is > 1 (i.e. *y* increases over time), while when the export probe is used, *k* corresponds to the export rate and *b* is < 1 (i.e. *y* decreases over time). Trust region reflective least-squares algorithm was used for the nonlinear regression. The fitting result was excluded from further analyses if it met one or more of the following criteria: (1) the fitting algorithm did not converge; (2) there were too many missing time points in the nuclear intensity trajectory (i.e. incomplete nucleus tracking); (3) reduced chi-squared statistics was too large; (4) |1-b| was too small; and (5) *k* was abnormally large.

The GFP images were corrected for camera dark noise and uneven illumination. Then, for each nucleus, mean GFP intensity was measured and converted to a concentration using the conversion factor obtained from the intensity calibration procedure described in Figure 1B and below. Dose-response curve was generated by plotting the nuclear transport rate (*k*) against the GFP concentration (*C*). The raw data points were logarithmically binned by the GFP concentration within the range between 0.1 μM and 40 μM. Median and interquartile range (IQR) were calculated in each bin and plotted as an error bar in the dose-response curve. Using the Levenberg-Marquardt least squares algorithm, the following Hill function was fitted to the binned median values:

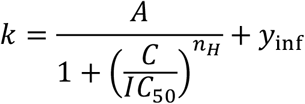

*A* is the amplitude of the inhibition (i.e., efficacy), *IC*_*50*_ the half-maximal inhibitory concentration (i.e., potency), and *n*_*H*_ the Hill coefficient.

#### GFP intensity calibration

The GFP intensity calibration (Figure 1B) was performed every 2-3 weeks or whenever there was a major change in the microscope (e.g., laser realignment). For the calibration, the U2OS stable cell line expressing H2A-Halo was transfected with the GFP-2A-ORF6 plasmid and prepared in an 8-well chamber slide in the same way as in the dose-response characterization. GFP and H2A-Halo:JF646 were imaged using the same microscope setting as in the dose-response characterization at ~10 different stage positions within a well. While the sample was on the stage, the recombinant GFP was added to the well at a certain concentration, and GFP and H2A-Halo:JF646 were imaged again at the same stage positions. This procedure was repeated at different wells with ~5 different concentrations of the added recombinant GFP, ranging from 0 to 5 μM. The GFP fluorescence intensities of each cell before and after adding the recombinant GFP (*I* and *I*_*in*_, respectively) were quantified in the same way as in the dose-response characterization. The extracellular background GFP intensity (*I*_*ex*_) in the presence the added recombinant GFP was determined by blurring the image with a gaussian filter with sigma = 2 pixels and finding the most frequent intensity value. For each recombinant GFP concentration (*C*), *I*_*in*_ – *I*_*ex*_ was plotted against *I*. A linear model was fitted to determine the x-intercept (*I*_*0*_). The slope of the linear model fitted to the plot of *I*_*0*_ against *C* was used as the intensity-to-concentration conversion factor. Levenberg-Marquardt least squares algorithm was used for the linear regressions.

### Western blot

U2OS cells were transfected with the GFP-2A-ORF6 or GFP-2A-ALFA-ORF6 plasmid via electroporation, using Nucleofector™ 2b device (Lonza) and Ingenio® electroporation kit (Mirus, #50117). After 1 day, the transfected cells were washed in ice-cold PBS, harvested via scrapping, resuspended in RIPA buffer (Thermo Fisher, #89900) supplemented with 1% SDS and protease inhibitor cocktail (Millipore Sigma, #11836170001), and briefly sonicated. The lysates were mixed with an equal volume of 2x SDS-PAGE sample buffer containing 100 mM DTT and heated at 95ºC for 5 min. Precast 4–12% gradient Bis-Tris gel (Thermo Fisher, #NP0321) was used for SDS-PAGE. Proteins were blotted onto a 0.2 μm nitrocellulose membrane (Bio-Rad, #1620213) via the standard wet electrotransfer protocol. The membrane was incubated in 50% methanol/water (v/v) for 30 min, followed by heating at 50ºC for 30 min. The fixed membrane was incubated in the Intercept Blocking Buffer (LI-COR, #927-70001) for 2 hours at RT and then in the blocking buffer containing monoclonal anti-GFP mouse antibody (GF28R, 1:3000, Thermo Fisher, #MA5-15256) and biotin-conjugated anti-ALFA nanobody (1:1000, NanoTag, N1505-Biotin) at 4ºC overnight. The membrane was washed 4 times in PBS with 0.1% Tween 20 and then incubated in the blocking buffer containing 1:20,000 IRDye^®^ 680RD goat anti-mouse IgG secondary antibody and IRDye^®^ 800CW streptavidin (LI-COR, #926-68070 and #926-32230) for 1 hour at RT. After being washed 4 times in PBS with 0.1% Tween 20 and once in PBS, the membrane was imaged using Odyssey DLx imaging system.

### Immunofluorescence

#### Sample preparation

U2OS cells were seeded at ~10,000 cells per well in a glass-bottom eight-well chambered coverslip (ibidi, #80827) with 250 μl complete DMEM. On the next day, cells in each well were transfected with a plasmid encoding GFP-2A-POI using TransIT-2020 transfection reagent as described above, where POI is an epitope tagged protein (e.g., ALFA-ORF6^CoV2^ and ORF6-ALFA^CoV2^). After ~24 hours of transfection, GFP images were acquired to determine the concentration of POI as described above.

Then, cells were fixed using 2% PFA/PBS for 20 min at RT, permeabilized using 25 μg/ml digitonin (Millipore Sigma, #300410) in PBS for 20 min at RT, and blocked using 1% BSA/PBS for 1 hour at RT. AF647-conjugated anti-ALFA nanobody (NanoTag, #N1502-AF647) and anti-NUP98 rabbit monoclonal antibody (C39A3, Cell Signaling, #2598) were diluted by factors of 500 and 50 in 1% BSA/PBS, respectively. Cells were incubated with the diluted antibodies overnight at 4°C in a humidified chamber. Then, cells were incubated with goat anti-rabbit secondary antibody conjugated with AF568 (Thermo Fisher Scientific, #A11036) diluted (1:1,000 ratio) in 1% BSA/PBS for 1 hour at RT. Cells were washed in PBS for 5 min three times after each incubation step.

#### Image acquisition & visualization

Immunofluorescence images were acquired using a Nikon Ti motorized inverted microscope equipped with Perfect Focus System, Yokagawa CSU-X1 spinning disk confocal, Nikon LUN-F XL solid state laser combiner, Hamamatsu ORCA-Fusion BT CMOS camera (6.5 μm^2^ photodiode), and Prior Proscan III motorized stage and shutters. CFI Plan Apo Lambda 100×/1.45-NA oil objective lens (Nikon) and Di01-T405/488/568/647 polychroic mirror (Semrock) and 200 ms exposure time were commonly used for all channels. 488-nm laser and ET525/50m filter (Chroma) were used for GFP imaging; 561-nm laser and ET605/52m filter (Chroma) for AF568; and 640-nm laser and ET705/72m filter (Chroma) for AF647. NIS-Elements software was used to control the hardware. OMERO.figure was used for image visualization. The same lookup table (LUT) was used for the same channel and the same construct, while the minimum was commonly set to the dark current noise of the camera (=100).

### Statistical information

All reported uncertainties of the parameter estimates are the standard errors. Data distribution was assumed to be normal in the uncertainty estimations, but this was not formally tested. Each dose-response curve consists of data acquired on at least 2 different days, and the total number of cells per dose-response curve is reported in figure legend. Batch effect was closely examined, but not detected.

## Supplemental information

Supplemental text: Rationale behind the use of fluorescence contrast for GFP intensity calibration

Figure S1. Example of GFP calibration data collected as illustrated in Figure 1B.

Figure S2. Negative control dose-response data.

Figure S3. Import vs export rates at different concentrations of ORF6.

Figure S4. Dose-response curve of ALFA-ORF6^CoV1^.

Figure S5. Dose-response curves obtained using different fluorescent proteins: mCerulean and mTurquoise2.

Figure S6. Dose-response curves of ORF6 mutants (for Figure 3).

Figure S7. Dose-response characteristics of ORF6^CoV2^ N- or C-terminally tagged with ALFA-tag, Flag-tag or HA-tag.

