## Supplemental information for "Quantification of nuclear transport inhibition by SARS-CoV-2 ORF6 using a broadly applicable live-cell dose-response pipeline"

### Supplemental Text

#### Rationale behind the use of fluorescence contrast for GFP intensity calibration

Our strategy for calibrating the GFP fluorescence intensity for conversion to the intracellular GFP concentration is based on finding the GFP intensities that camouflage the cells in the presence of recombinant GFP added to the culture at known concentrations. The key underlying assumption is simple: a cell is brighter than the surrounding background if the intracellular GFP concentration is higher than the extracellular GFP concentration and vice versa. The molecular brightness of GFP has previously been measured to be identical inside HeLa cells and in aqueous solution, supporting the assumption[1]. However, the use of recombinant GFP as a concentration reference may result in some caveats. The concentration of the recombinant GFP stock solution was determined using the absorbance at 488 nm and the previously reported molecular extinction coefficient value (53,300 $M^{-1}cm^{-1}$)[2]. The reported extinction coefficient varies across different sources by <10%, which may introduce a systematic error to the calibration measurement. Immature non-fluorescent GFP may exist in the cell and in the recombinant GFP stock solution to different extents, which may also introduce an error.

This approach, however, has advantages outweighing the caveats. The biggest advantage is that it is broadly applicable to any type of fluorescence microscopy without the need for a specialized instrument. As long as the fluorescence contrast between the inside and outside regions of a cell can be measured, microscopic settings such as the confocality (confocal vs wide-field), excitation photons (singe-photon vs multi-photon), lateral/axial resolutions, and magnification do not affect the applicability. Therefore, it can be easily combined with any other microscopy-based assays to correlate the absolute GFP concentration with other measurable phenotypes. As in this study, when applied to a low-magnification microscopy setting, this approach can measure the absolute concentration of a large number of single cells in a scalable manner.

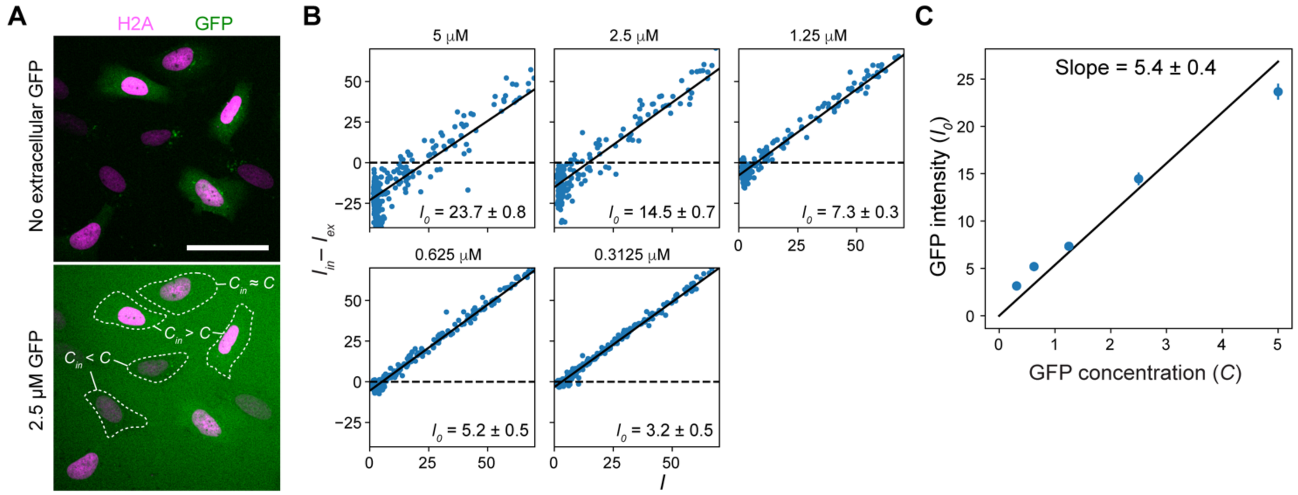


**Figure S1. Example of GFP calibration data collected as illustrated in Figure 1B.**
(A) (Steps 1 and 2) Representative images of U2OS cells transiently transfected with a GFP-expressing plasmid (top) before and (bottom) after adding 2.5 µM recombinant GFP. H2A-Halo was stably expressed, stained with JF646 and used as a nuclear marker. $C_{in}$ and $C$ denote the intracellular and extracellular GFP concentrations, respectively. 50 µm scale bar.
(B) (Step 3) Example plots of the difference between the intracellular and extracellular GFP intensities ($I_{in} - I_{ex}$) measured in the presence of a known concentration ($C$) of recombinant GFP in the culture vs the intracellular GFP intensity measured prior to the addition of the recombinant GFP ($I$). The black line represents the best line fit, and $I_0$ is the x-intercept, which corresponds to the GFP intensity of a cell whose intracellular GFP concentration is the same as the concentration of recombinant GFP added to the culture.
(C) Example plot of $I_0$ vs $C$. The black line represents the best line fit, of which slope serves as the intensity-to-concentration conversion factor.

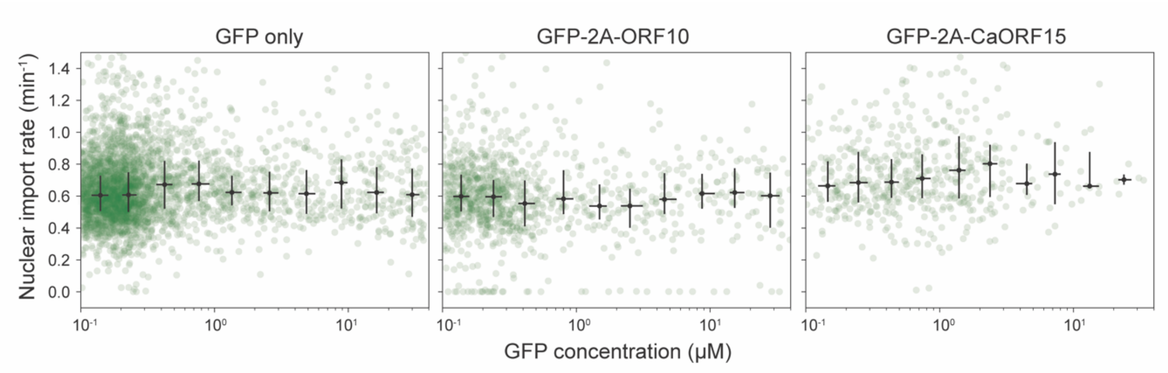


**Figure S2. Negative control dose-response data.**

Dose-response curves for (left) GFP expression only and for co-expression of GFP and (middle) ORF10 or (right) candidate ORF15 of SARS-CoV-2. n>500 cells for each. Green circles represent raw data, and black error bars represent median and interquartile ranges in each bin.

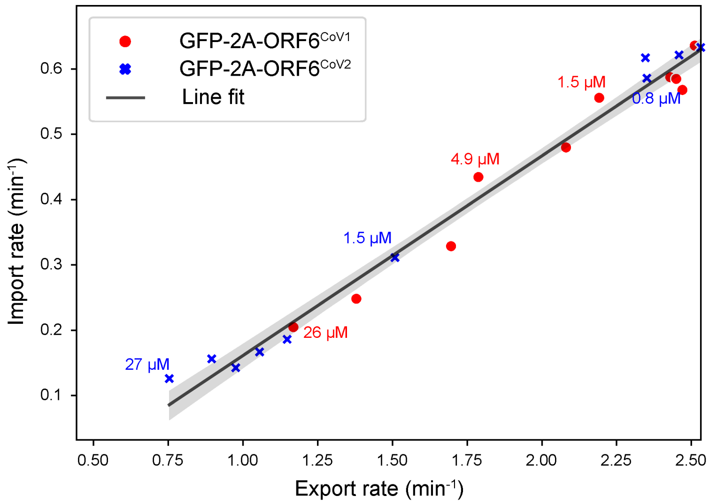


**Figure S3. Import vs export rates at different concentrations of ORF6.**
The data points are the medians binned by GFP concentration shown in the dose-response curves in Figure 2B. The black line represents the line fitted to the combined data of $ORF6^{CoV1}$ (red circle) and $ORF6^{CoV2}$ (blue x), and the black shaded area is the corresponding 95% confidence interval.

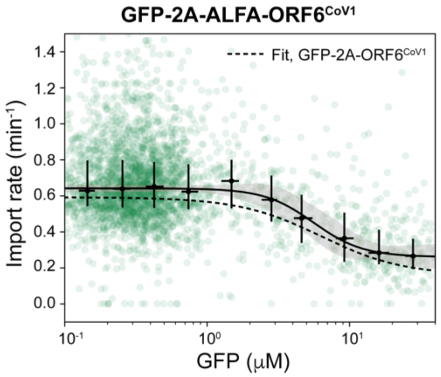


**Figure S4. Dose response curves for the nuclear import inhibition by ALFA-ORF6$^{CoV1}$**. n>1000 cells. The dashed line is the Hill function fit for the untagged ORF6$^{CoV1}$ for comparison. The green circles are raw data points, and the black error bars are medians and interquartile ranges binned by GFP concentration. Black line represents the best fitted Hill function, and black shaded area is the corresponding 95% confidence interval.

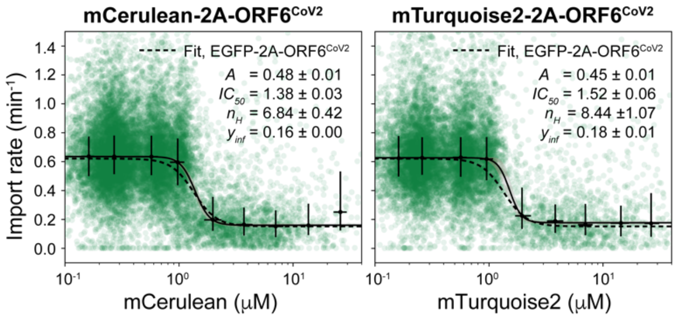


**Figure S5. Dose-response curves obtained using different fluorescent proteins.** mCerulean-2A-ORF6$^{CoV2}$ (left) and mTurquoise2-2A-ORF6$^{CoV2}$ (right). The dashed line is the Hill function fit for the EGFP-2A-ORF6$^{CoV2}$ for comparison. The green circles are raw data points, and the black error bars are medians and interquartile ranges binned by fluorescent protein concentration. Black line represents the best fitted Hill function, and black shaded area is the corresponding 95% confidence interval.

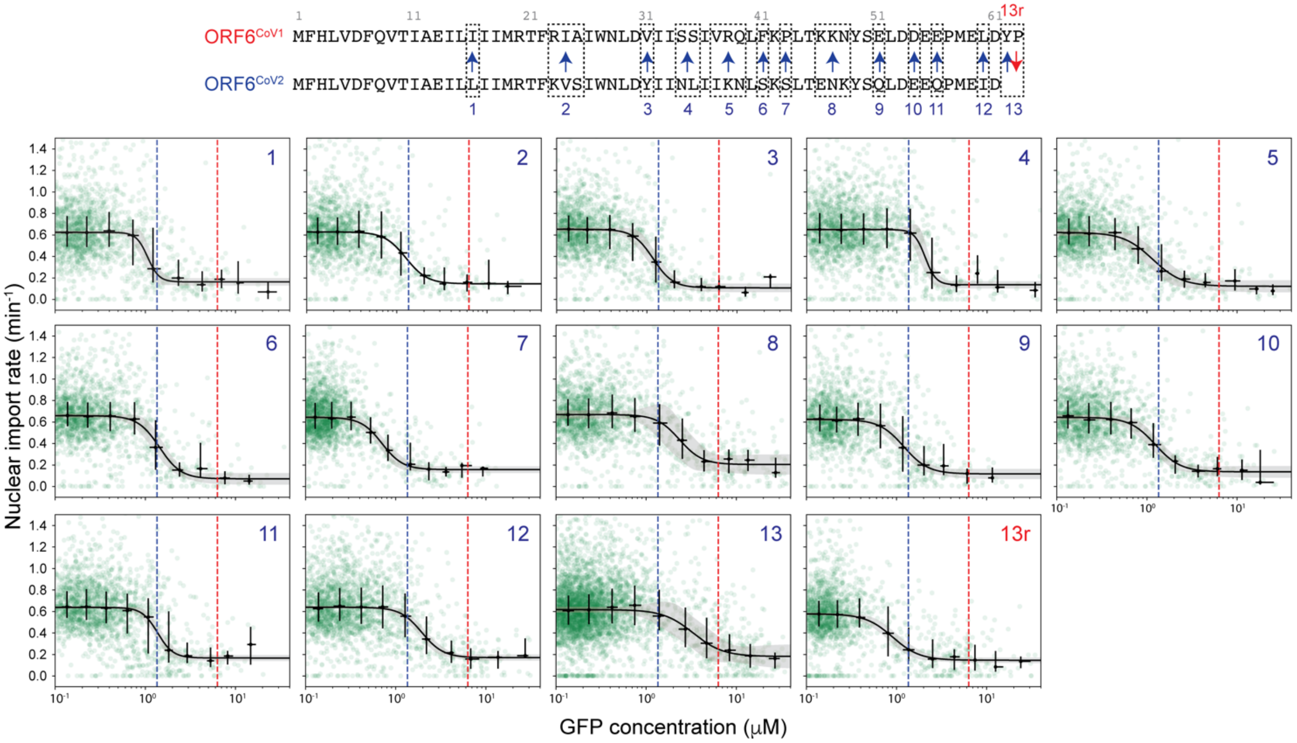

**Figure S6. Dose-response curves of ORF6 mutants (for Figure 3).** n>1300 cells for each. The green circles are raw data points, and the black error bars are medians and interquartile ranges binned by GFP concentration. Black line represents the best fitted Hill function, and black shaded area is the corresponding 95% confidence interval. The red and blue vertical dashed lines indicate the $IC_{50}^{GFP}$s of the ORF6$^{CoV1}$ and ORF6$^{CoV2}$ wild types, respectively.

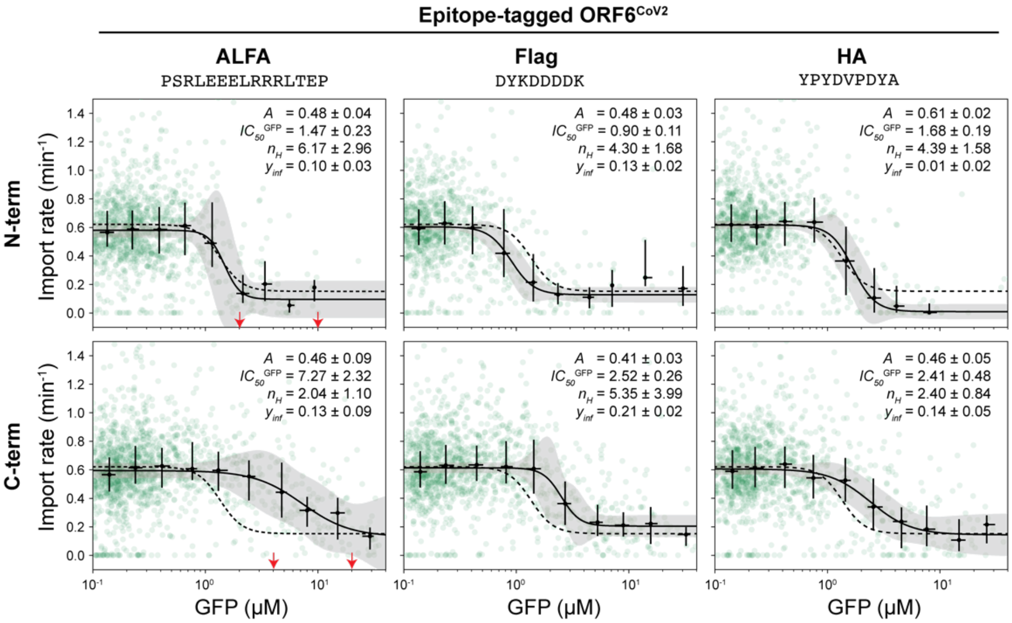


**Figure S7. Dose-response characteristics of ORF6$^{CoV2}$ N- or C-terminally tagged with ALFA-tag, Flag-tag, or HA-tag.**

n > 850 cells for each construct. The green circles are raw data points, and the black error bars are medians and interquartile ranges binned by GFP concentration. The black solid line represents the best fitted Hill function, and the black shaded area is the corresponding 95% confidence interval. The black dashed line is the Hill function fit for the ORF6$^{CoV2}$ wild type shown for comparison. The plot for the ALFA-ORF6$^{CoV2}$ (upper left) is the same as Figure 2C but placed here again to facilitate comparison. Red arrows indicate the GFP concentrations of the cells shown in the immunofluorescence images in Figure 5A and B.
